# SBGNview: Data Analysis, Integration and Visualization on All Pathways

**DOI:** 10.1101/2021.03.26.437200

**Authors:** Xiaoxi Dong, Kovidh Vegesna, Cory Brouwer, Weijun Luo

## Abstract

Pathway analysis is widely used in genomics and omics research, but the data visualization has been highly limited in function, pathway coverage and data format. Here we develop SBGNview a comprehensive solution to address these needs. By adopting the standard SBGN format, SBGNview greatly extend the coverage of pathway based analysis and data visualization to essentially all major pathway databases beyond KEGG, including 5200 reference pathways and over 3000 species. In addition, SBGNview substantially extends current tools in both design and function, including standard coherent input/output formats, high quality graphics convenient for both computational and manual analysis, and flexible and open-end workflow. In addition to pathway analysis and data visualization, SBGNview provides essential infrastructure for SBGN data manipulation and processing. SBGNview is available online: https://github.com/datapplab/SBGNview.

## Introduction

Pathway analysis has become a prevalent analytical approach in genomics and omics studies. Numerous analysis methods^1,2^ and databases^3-8^ have been developed. But tools for pathway analysis visualization are sparse, with Pathview^9,10^ and Cytoscape^11^ as two representatives. Cytoscape renders pathways as networks efficiently, which tend to lose the context information and are hard to interpret. Pathview^9,10^, a tool we developed previously, can map various biological data and generate highly interpretable pathway graphs with biological context, but only supports KEGG^3^ and its pathway data format (KGML). Major pathway databases like KEGG^12^ and Reactome^13^ also provide integrated yet limited functions for data analysis and visualization. There has not been a tool for data integration and visualization supporting pathways cross major databases and standard formats.

Here we present a new tool set --SBGNview as a systematic solution to this pressing need. SBGNview is built on Systems Biology Graphical Notation (SBGN)^14^, a set of high quality, standard graphical languages for representing biological processes and interactions. SBGN has been widely adopted and supported by major pathway databases, collections, and pathway curation or editing packages (https://SBGN.github.io/software). Unfortunately, SBGN pathways so far have limited usage in data integration and visualization. We developed SBGNview to fill in this gap. Consequentially, all major pathway databases adopting SBGN format are now open to pathway analysis and data visualization in a consistent and robust way, including Reactome^4^, PANTHER^5^, SMPDB^6^, MetaCyc^7^, MetaCrop^15^ and Pathway Commons^8^. This is much broader and deeper collection of reference pathways than KEGG alone, in terms of pathway number, categories, resolution and details. In addition to functions in data integration and visualization, SBGNview provides a comprehensive tool set for SBGN based pathway analysis, as well as SBGN data search, retrieval, and processing.

## Main Features

### SBGNview greatly extends the coverage of pathway based analysis and data visualization

SBGNview naturally support all pathway databases and collections that adopt SBGN standards and its format (SBGN-ML)^16,17^, and make them accessible for pathway analysis and data visualization (Table 1 and Supplementary Table 1). SBGNview provides two tiers of support for SBGN pathway data. Tier 1, deep and direct support to a core collection of pathway data from 5 major pathway databases including Reactome^4^, PANTHER^5^, SMPDB^6^, MetaCyc^7^ and MetaCrop^15^. These databases together covers 5,200 reference pathways and over 3,000 species (Table 1). This is a much broader and deeper collection of reference pathways than KEGG alone, especially in the domains of major research species, crops, small molecules, and metabolic pathways. In addition to the 3,000 species from the original databases, SBGNview can also map reference pathways to 6,190 KEGG species via KEGG Orthology^18^. SBGNview provides diagram optimization and ID mapping on these pathway databases. In other words, we’ve prepared a full collection of high quality SBGN pathways from these databases, which can be directly used in data analysis. The original pathway diagram/layout have been extensively improved, and nodes, edges and other graph components extensively adjusted in sizes and position as to remove, reduce or balance the overlap and the blank space between them (Fig. 1 vs Supplementary Fig. 2). These improvements are essential for data visualization, human interpretation and publication purpose. In addition, glyph IDs of the original SBGN-ML files are largely databases specific, not common molecule IDs. We generated ID mapping between SBGN-ML glyph IDs and common molecule IDs, so that omics or molecular data can be easily mapped to these pathways. Tier 2, generic support to all other pathway databases, collections and users’ custom pathway data in SBGN format (Supplementary Table 1). These data can be used in SBGNview, except diagrams may not be optimized for data visualization, and users need to provided ID mapping or make sure the glyph IDs are commonly use molecule IDs. Note the collection of pathways in databases with Tier 1 support is comprehensive, and should cover most of the user needs in pathway analysis and data visualization.

**Table 1.** SBGN pathway databases directly supported by SBGNview (Tier 1 support).

| Database | SBGN data | Pathways<sup>*</sup> | Species | Description |
| --- | --- | --- | --- | --- |
| Reactome | Pathway Commons | 1,746 | 16 | Pathway database for major research species |
| SMPDB | Pathway Commons | 725 | 1 | Human small molecule pathways |
| PANTHER | Pathway Commons | 149 | 132 | Signaling pathways of multiple species |
| MetaCyc | MetaCyc | 2,518 | 2,980 | Metabolic pathways from all domains of life |
| MetaCrop | MetaCrop | 62 | 9 | Crop metabolism pathways |
| KEGG<sup>#</sup> | KEGG | 430 | 6,190 | Pathway database for most sequenced species |

<sup>\*</sup>Ortholog pathways across multiple species counted as the same pathway, similar to reference pathway in KEGG. <sup>#</sup>KEGG statistics for comparison. All data were collected in 2020.

**Figure 1.**
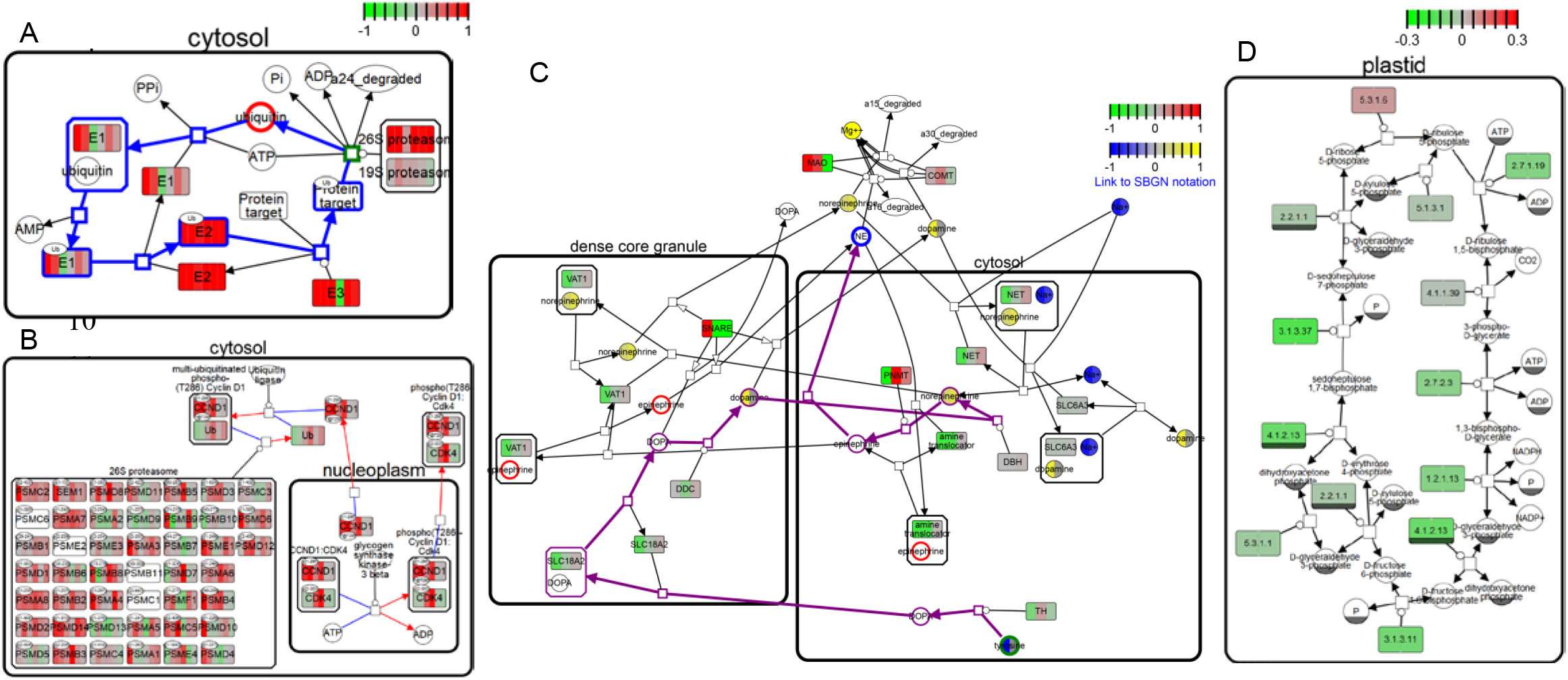
Example SBGNview graphs: A) Ubiquitin proteasome pathway (PANTHER pathway P00060) and B) Ubiquitin-dependent degradation of Cyclin D1 pathway (Reactome pathway R-HSA-69229) up-regulated in breast cancer (GEO GSE16873); C) both gene data (GEO GSE16873) and compound data (simulated with Pathview) with highlighted nodes and path on the Adrenaline and noradrenaline biosynthesis pathway (PANTHER pathway P00001); D) Calvin cycle (MetaCrop pathway) downregulated in Arabidopsis pen3 mutants (GEO GSE3220). In C), Nodes highlighted in red are epinephrine, the shortest path between tyrosine (start node in green) and epinephrine (end node in blue) is highlighted in purple.

### SBGNview extends the architecture and workflow of Pathview with numerous unique features

SBGNview has a similar design as the Pathview package^9^ in four functional modules: Downloader, Parser, Mapper and Viewer (Supplementary Fig. 1). These modules are the integral components of SBGNview, but they are also useful for SBGN data input/output, processing, parsing and rendering in general. In addition to these Pathview-like modules, SBGNview workflow has a unique part, the Highlighter module. Highlighter provides a post-rendering modification mechanism, which can highlight or modify any part(s) of the initial rendering of the SBGN graph object (Fig. 1c and Supplementary Fig. 3). The updated rendering then can then be re-fed into Viewer for updated graphics output. In other words, the Highlighter makes SBGNview workflow an open-ended process, suitable for iterative editing/updating and interactive visualization.

SBGNview also differs from Pathview^9^ in both input and output. For input pathway data, SBGNview supports the standard SBGN format adopted by multiple major databases, while Pathview supports KGML used only by KEGG (Supplementary Table 2). Pathview has two discrete output styles, KEGG view and Graphviz view. The former is a raster image with all context information (tissue/cell types etc) labelled by KEGG curator. The latter is a vector image with no context information. SBGNview has one unified output style (with different file formats, i.e. SVG, PDF, PNG and PS), a high resolution vector image with comprehensive meta-data and context information defined with SBGN-ML. Even more importantly, the primary output format SVG, as XML based graphics, is readable by both machine and human, and accessible for either automatic or manual updating/editing.

### SBGNview provides a comprehensive tool set for SBGN pathway analysis, data processing and visualization

SBGNview integrates 3 primary functions in working with SBGN pathways: data visualization, data integration and pathway analysis workflow. For Data visualization, SBGNview offers extensive graphics control on all elements (glyphs, complex, compartments, arcs, reactions, processes, text, etc) and their attributes (line type, width, shape, strike, fill, color, labels, size, position, even margins etc) in the pathway diagrams. The Highlighter module in SBGNview provides extra mechanism of graphics control, targeting specific graphic attributes of selected sets, subsets or classes of pathway components (proteins/enzymes, interactions, reactions, processes) or graph elements (glyphs, arcs and paths) (Supplementary Fig. 3). This is especially useful for sub-pathway level analysis, visualization and interpretation. For data integration, by connecting to Bioconductor^19^ and KEGG^3^ annotation resources, SBGNview can maps, integrates and renders a large variety of biological data on SBGN pathways. It supports most common gene or compound ID types, thousands of reference pathways and species (Table 1). Like Pathview, SBGNview can be easily integrated with a wide variety of existing tools for omics data analysis and pathway analysis. In one package vignette, we demonstrated a complete pathway analysis workflow using GAGE^1^ + SBGNview.

SBGNview also provides general infrastructure for SBGN pathway data processing. KEGG^3^ hosts a centralized REST API for pathway data search, retrieval, extraction and downloading. Pathview takes the advantage of the KEGG API for these tasks. Unfortunately, there has been no such facility for SBGN pathway data largely because the data come from many heterogeneous sources. SBGNview fills in this gap as the first tool focusing on SBGN pathway based data analysis. The 5 functional modules of SBGNview are general useful for SBGN pathway data processing hence an important infrastructure. In addition, we provide a high quality collection of SBGN pathway data with open access via GitHub (SBGNhub: https://github.com/datapplab/SBGNhub). We also provide functions, data and supportive package (SBGNview.data: https://github.com/datapplab/SBGNview.data) for SBGN pathway search, retrieval, data mapping and extraction. For pathway analysis, SBGNview has functions for pathway gene sets extraction and ID conversion.

## Conclusion

SBGNview maps, integrates and renders a wide range of biological data to SBGN pathways. By adopting the standard SBGN format, SBGNview greatly extend the coverage of pathway based analysis and data visualization to essentially all major pathways beyond KEGG. SBGNview extends the proven design of Pathview with multiple unique features, including standard and widely supported input and output formats, high resolution graphics accessible to both machine and human, and flexible, iterative and open-ended workflow. While Pathview primarily focuses on data visualization, SBGNview provides a complete workflow and comprehensive tool set for SBGN based pathway analysis, data visualization and processing.

## Data Availability

The SBNGview package provides the ready-to-use SBGN pathway data collection with Tier 1 support. Additional pathway data, demo and supportive datasets are included in SBGNview, SBGNview.data and SBGNhub packages. Please see the package documentation and tutorials for details.

We used a human breast cancer microarray dataset as the demo gene data. The raw data is accessible in GEO (GSE16873). The processed data is provided as part of SBGNview package. This dataset was used in Fig 1A-C and Supplementary Fig. 3-6. The Arabidopsis pen3 mutant dataset was used in Fig. 1D. Both the raw and processed data were retrieved from GEO (GSE3220). The compound dataset used in Fig 1C and Supplementary Fig. 3 was simulated using Pathview, with code available in the main tutorial.

## Code Availability

SBGNview package is available on GitHub: https://github.com/datapplab/SBGNview. We also provide supportive data and additional functions in the SBGNview.data package: https://github.com/datapplab/SBGNview.data.

## References

1 Luo, W., Friedman, M. S., Shedden, K., Hankenson, K. D. & Woolf, P. J. GAGE: generally applicable gene set enrichment for pathway analysis. BMC Bioinformatics 10, 161 (2009).

2 Khatri, P., Sirota, M. & Butte, A. J. Ten years of pathway analysis: current approaches and outstanding challenges. PLoS Comput Biol 8, e1002375, doi:10.1371/journal.pcbi.1002375 (2012).

3 Kanehisa, M., Furumichi, M., Tanabe, M., Sato, Y. & Morishima, K. KEGG: new perspectives on genomes, pathways, diseases and drugs. Nucleic acids research 45, D353–D361, doi:10.1093/nar/gkw1092 (2017).

4 Jassal, B. et al. The reactome pathway knowledgebase. Nucleic Acids Res 48, D498–D503, doi:10.1093/nar/gkz1031 (2020).

5 Mi, H. et al. PANTHER version 11: expanded annotation data from Gene Ontology and Reactome pathways, and data analysis tool enhancements. Nucleic Acids Res 45, D183–D189, doi:10.1093/nar/gkw1138 (2017).

6 Jewison, T. et al. SMPDB 2.0: big improvements to the Small Molecule Pathway Database. Nucleic Acids Res 42, D478–484, doi:10.1093/nar/gkt1067 (2014).

7 Caspi, R. et al. The MetaCyc database of metabolic pathways and enzymes - a 2019 update. Nucleic Acids Res 48, D445–D453, doi:10.1093/nar/gkz862 (2020).

8 Rodchenkov, I. et al. Pathway Commons 2019 Update: integration, analysis and exploration of pathway data. Nucleic Acids Res 48, D489–D497, doi:10.1093/nar/gkz946 (2020).

9 Luo, W. & Brouwer, C. Pathview: an R/Bioconductor package for pathway-based data integration and visualization. Bioinformatics 29, 1830–1831, doi:10.1093/bioinformatics/btt285 (2013).

10 Luo, W., Pant, G., Bhavnasi, Y. K., Blanchard, S. G., Jr. & Brouwer, C. Pathview Web: user friendly pathway visualization and data integration. Nucleic Acids Res, doi:10.1093/nar/gkx372 (2017).

11 Smoot, M. E., Ono, K., Ruscheinski, J., Wang, P. L. & Ideker, T. Cytoscape 2.8: new features for data integration and network visualization. Bioinformatics 27, 431–432, doi:10.1093/bioinformatics/btq675 (2011).

12 Kanehisa, M. & Sato, Y. KEGG Mapper for inferring cellular functions from protein sequences. Protein Sci. 29, 28–35, doi:10.1002/pro.3711 (2020).

13 Sidiropoulos, K. et al. Reactome enhanced pathway visualization. Bioinformatics 33, 3461–3467, doi:10.1093/bioinformatics/btx441 (2017).

14 Le Novere, N. et al. The Systems Biology Graphical Notation. Nat. Biotechnol. 27, 735–741, doi:10.1038/nbt.1558 (2009).

15 Schreiber, F. et al. MetaCrop 2.0: managing and exploring information about crop plant metabolism. Nucleic Acids Res 40, D1173–1177, doi:10.1093/nar/gkr1004 (2012).

16 van Iersel, M. P. et al. Software support for SBGN maps: SBGN-ML and LibSBGN. Bioinformatics 28, 2016–2021, doi:10.1093/bioinformatics/bts270 (2012).

17 Rougny, A. et al. Systems Biology Graphical Notation: Process Description language Level 1 Version 2.0. Journal of integrative bioinformatics 16, doi:10.1515/jib-2019-0022 (2019).

18 Kanehisa, M., Sato, Y., Kawashima, M., Furumichi, M. & Tanabe, M. KEGG as a reference resource for gene and protein annotation. Nucleic acids research 44, D457–462, doi:10.1093/nar/gkv1070 (2016).

19 Gentleman, R. C. et al. Bioconductor: open software development for computational biology and bioinformatics. Genome Biol 5, R80 (2004).

