## Supplementary Notes, Figures & Tables for "SBGNview: Data Analysis, Integration and Visualization on All Pathways"

**This PDF file includes:**

**Example analysis**

**Supplementary Table 1-2**
**Supplementary Table 3-4 Captions**
**Supplementary Figs. 1-6**

### Example analysis

We illustrate SBGNview's functions in pathway analysis and data visualization using a human breast cancer microarray dataset (GSE16873)[1]. We downloaded the microarray raw data from GEO and process the gene expression data using RMA[2], collected gene sets from SBGN pathways using SBGNview, performed pathway enrichment analysis using GAGE[3], and mapping and rendered user data with significant pathways using SBGNview.

The GSE16873 dataset covers twelve patient cases, each with HN (histologically normal), ADH (ductal hyperplasia), and DCIS (ductal carcinoma in situ) RNA samples. For this example analysis, we compared DCIS to HN samples in the first 6 patients. We did two pathway analyses, one with the SBGNhub pathway collection of SBGNview, another with KEGG pathways. Note the SBGNhub collection include pathways from Reactome, PANTHER, SMPDB, MetaCyc, and MetaCrop. We identified 125 up-regulated and 26 down-regulated pathways in the SBGNhub collection (Supplementary Table 3), 23 up and 6 down regulated pathways from KEGG (Supplementary Table 4). Indeed, SBGNhub represents a much more comprehensive coverage of pathways than KEGG.

Overall, there is a high level of consistence between the two analyses. Most significant KEGG pathways can found similar entries in the SBGNhub list, but the latter is much more inclusive, specific and informative. For example, there is one cell cycle pathway in KEGG (hsa04110 Cell cycle), but numerous in the SBGNhub list, including R-HSA-2467813::Separation of Sister Chromatids, R-HSA-174184::Cdc20:Phospho-APC/C mediated degradation of Cyclin A, R-HSA-8852276::The role of GTSE1 in G2/M progression after G2 checkpoint and many more.

Importantly, we discovered numerous pathways in the SBGNhub pathway analysis that were missed by the KEGG pathway analysis. For instance, DNA damage repair (Reactome: R-HSA-5693565), NF-kB signaling (Reactome: R-HSA-5676590) and apoptosis (PANTHER: P00006) pathways were dis-regulated in breast cancer samples based on SBGNhub pathway analysis (Supplementary Table 3), but not KEGG pathway analysis (Supplementary Table 4).

Visualization of fold changes between in cancer vs normal samples revealed relevant details of the perturbation. In Ubiquitin-dependent degradation of Cyclin D1 pathway (Reactome pathway R-HSA-69229) (Fig. 1B), some members (e.g. PSMA7) of the 26S proteasome are up-regulated much more than other members (e.g. PSMD9) in the same complex. On the other hand, genes are unanimously or extensively down-regulated in Non-integrin membrane-ECM interaction (Reactome pathway R-HSA-3000171), Smooth Muscle Contraction (Reactome pathway R-HSA-445355) and Nicotinic acetylcholine receptor signaling pathway (PANTHER pathway P00044) (Supplementary Fig. 4-6).

### References


1 Emery, L. A. *et al.* Early dysregulation of cell adhesion and extracellular matrix pathways in breast cancer progression. *Am. J. Pathol.* **175**, 1292-1302, doi:10.2353/ajpath.2009.090115 (2009).

2 Irizarry, R. A. *et al.* Exploration, normalization, and summaries of high density oligonucleotide array probe level data. *Biostatistics* **4**, 249-264 (2003).

3 Luo, W., Friedman, M. S., Shedden, K., Hankenson, K. D. & Woolf, P. J. GAGE: generally applicable gene set enrichment for pathway analysis. *BMC Bioinformatics* **10**, 161 (2009).

| **Database** | **Pathways*** | **Species** | **Description** |
|---|---|---|---|
| ACSN | 15 | human | Cancer Signalling |
| AlzPathway | 1 | human | Alzheimer's disease |
| iPAVS | 500+ | human | manually curated human pathways |
| NaviCell | 20 | mammals; yeast | Signaling and disease pathways |
| Parkinson’s disease map | 1 | human | Parkinson’s disease |

Supplementary Table 1. Example SBGN pathway databases indirectly supported by SBGNview (Tier 2 support). *Ortholog pathways across multiple species counted as the same pathway, similar to reference pathway in KEGG. All data were collected in 2020.

| Format | KEGG | SBGN |
|---|---|---|
| Language | KGML | SBGN-PD |
| Entity/node | entry | glyph |
| Connection/edge | relation | arc |
| Step | reaction | process glyph |
| Complex container | group entry | complex glyph |
| Compartment container | none | compartment glyph |
| Image type | raster | vector |
| Resolution | low | high |
| Diagram plotting | semi-manual | automatic |
| Supported by | KEGG | most major databases |

Supplementary Table 2. Comparison between KEGG and SBGN pathway data formats. Note only Process Description (PD) is used, Activity Flow (AF) and Entity Relationship (ER) are not, because the former is the most SBGN common language used in pathway definition.

Supplementary Table 3 (separate Excel file). Significant sbgn pathways in DCIS vs HN comparison in GSE16873 dataset. Pathway analysis was done using GAGE (Luo et al, 2009).

Supplementary Table 4 (separate Excel file). Significant KEGG pathways in DCIS vs HN comparison in GSE16873 dataset. Pathway analysis was done using GAGE (Luo et al, 2009).

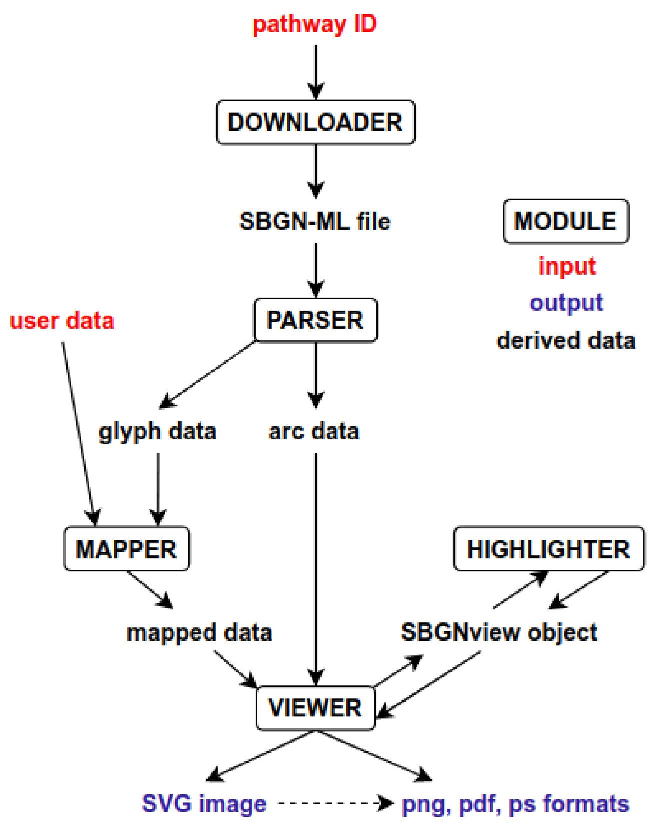


Supplementary Figure 1. SBGNview architecture and workflow.

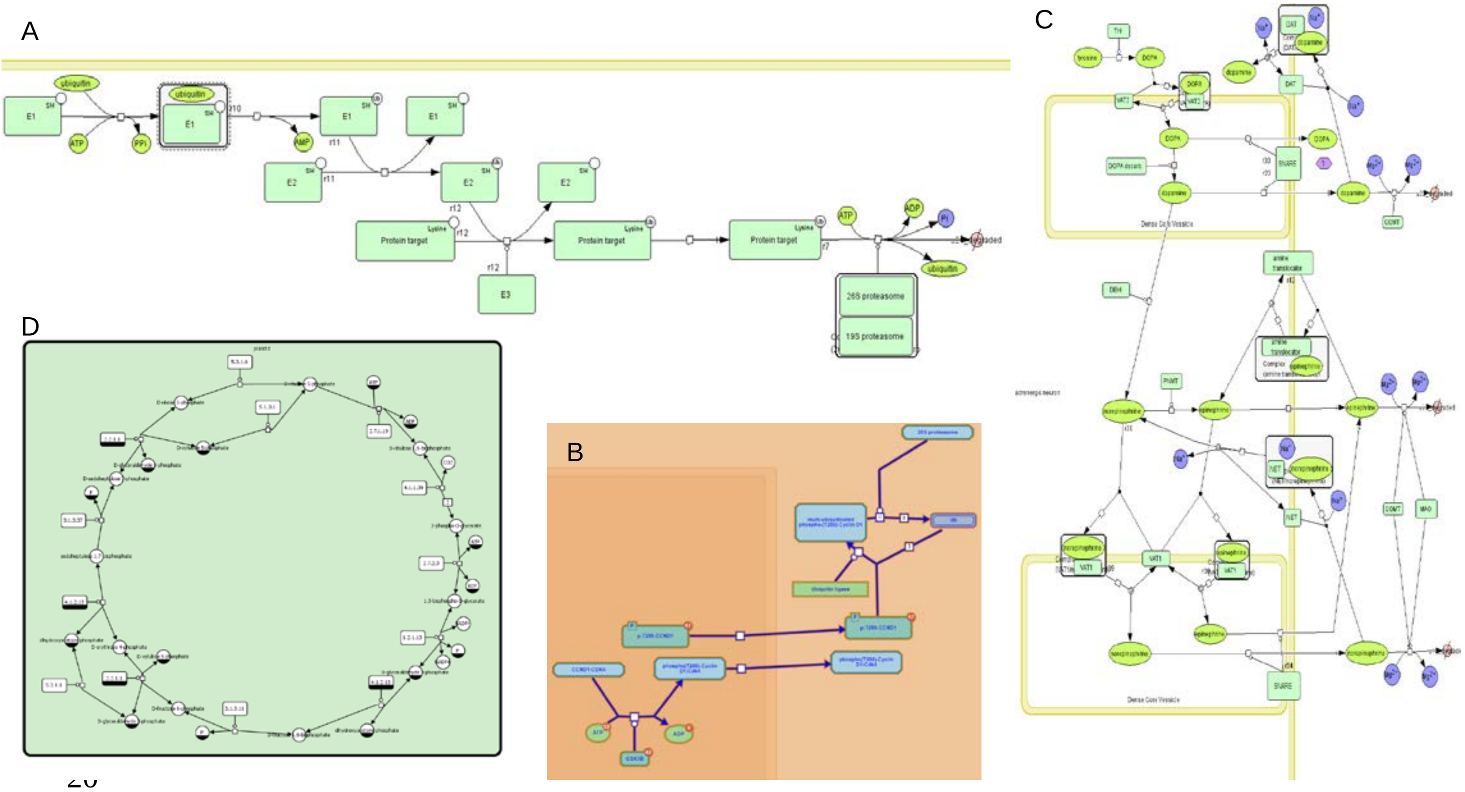


Supplementary Figure 2. Example Sbgn pathway diagrams in the original databases: A) Ubiquitin proteasome pathway (PANTHER pathway P00060) ; B) Ubiquitin-dependent degradation of Cyclin D1 pathway (Reactome pathway R-HSA-69229) ; C) Adrenaline and noradrenaline biosynthesis pathway (PANTHER pathway P00001); D) Calvin cycle (MetaCrop pathway).

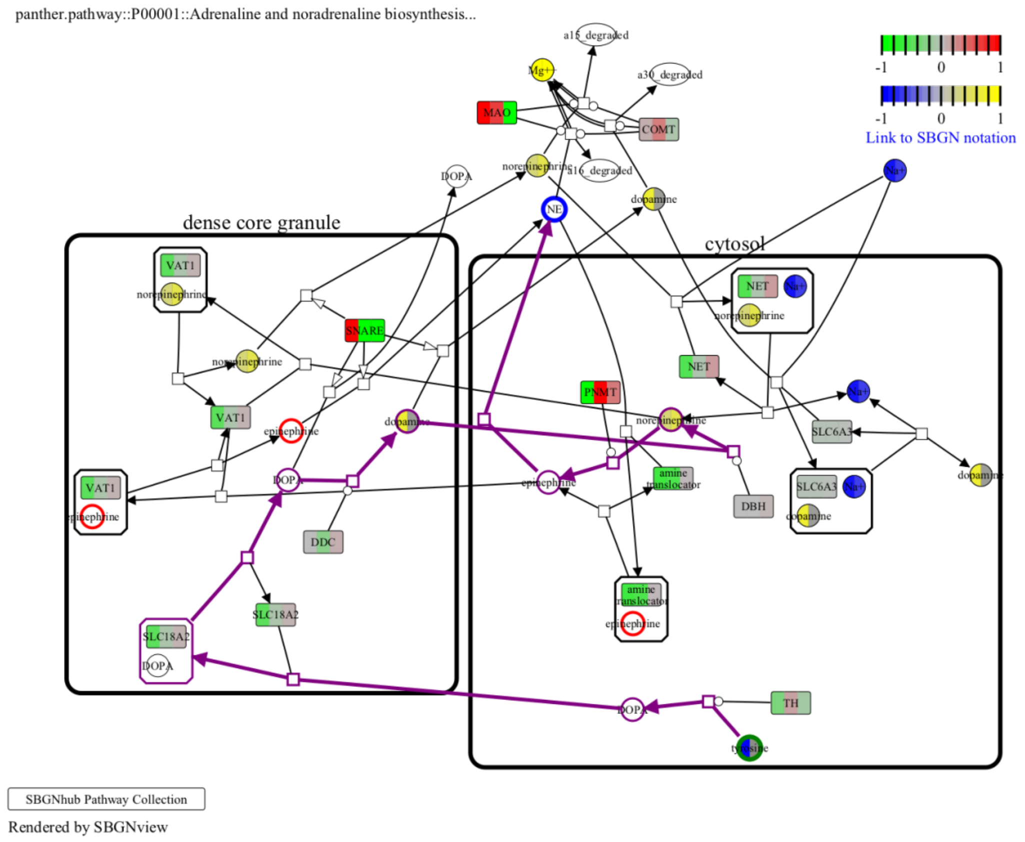


Supplementary Figure 3. Example SBGNview graph with both gene data (GEO GSE16873) and compound data (simulated with Pathview) and highlighted nodes and path on the Adrenaline and noradrenaline biosynthesis pathway (PANTHER pathway P00001). Nodes highlighted in red are epinephrine, the shortest path between tyrosine (start node in green) and epinephrine (end node in blue) is highlighted in purple.

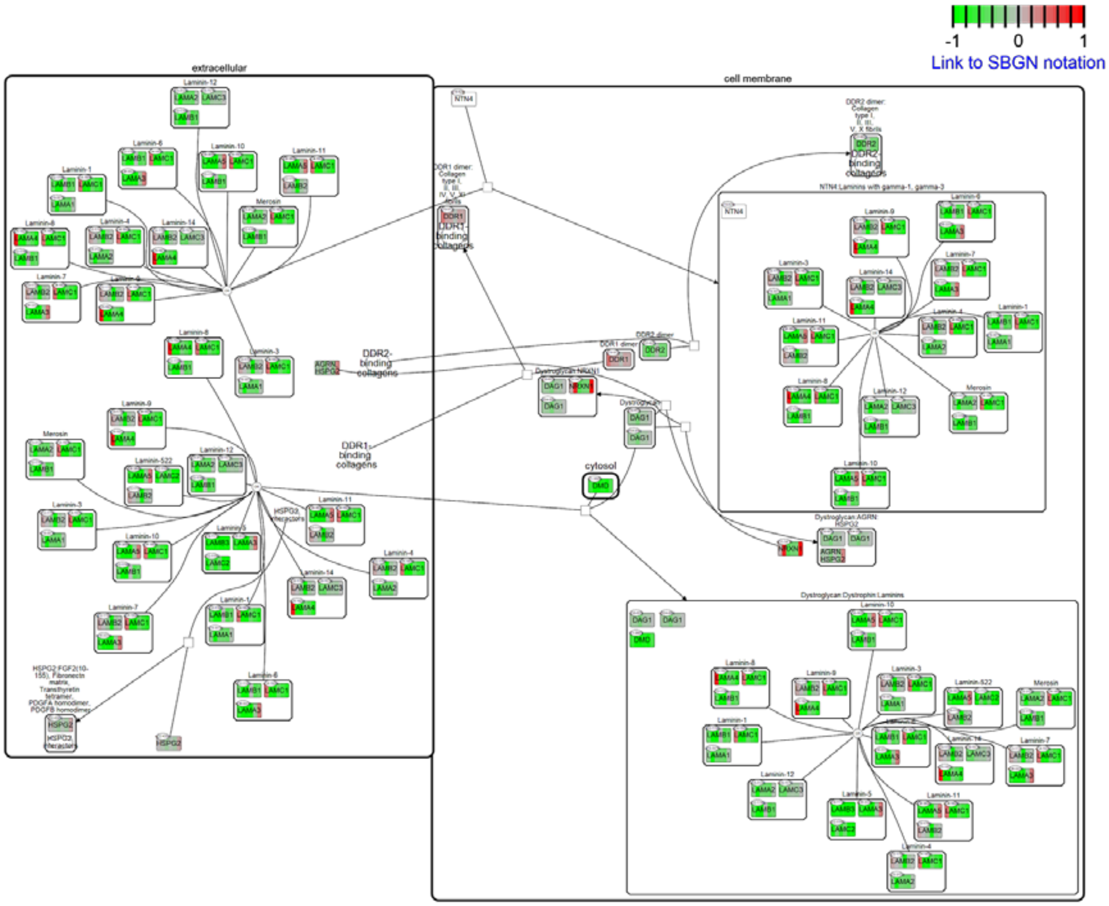


Supplementary Figure 4. Example SBGNview graph: Non-integrin membrane-ECM interaction (Reactome pathway R-HSA-3000171) down-regulated in breast cancer (GEO GSE16873).

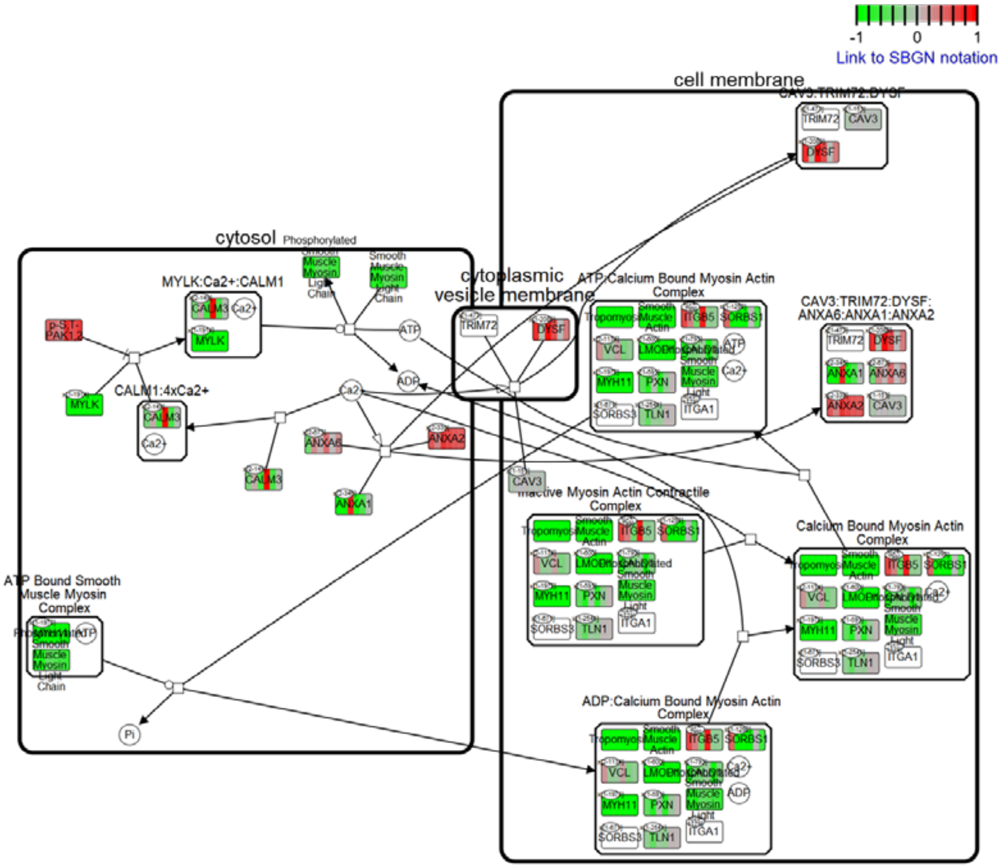


Supplementary Figure 5. Example SBGNview graph: Smooth Muscle Contraction (Reactome pathway R-HSA- 445355) down-regulated in breast cancer (GEO GSE16873).

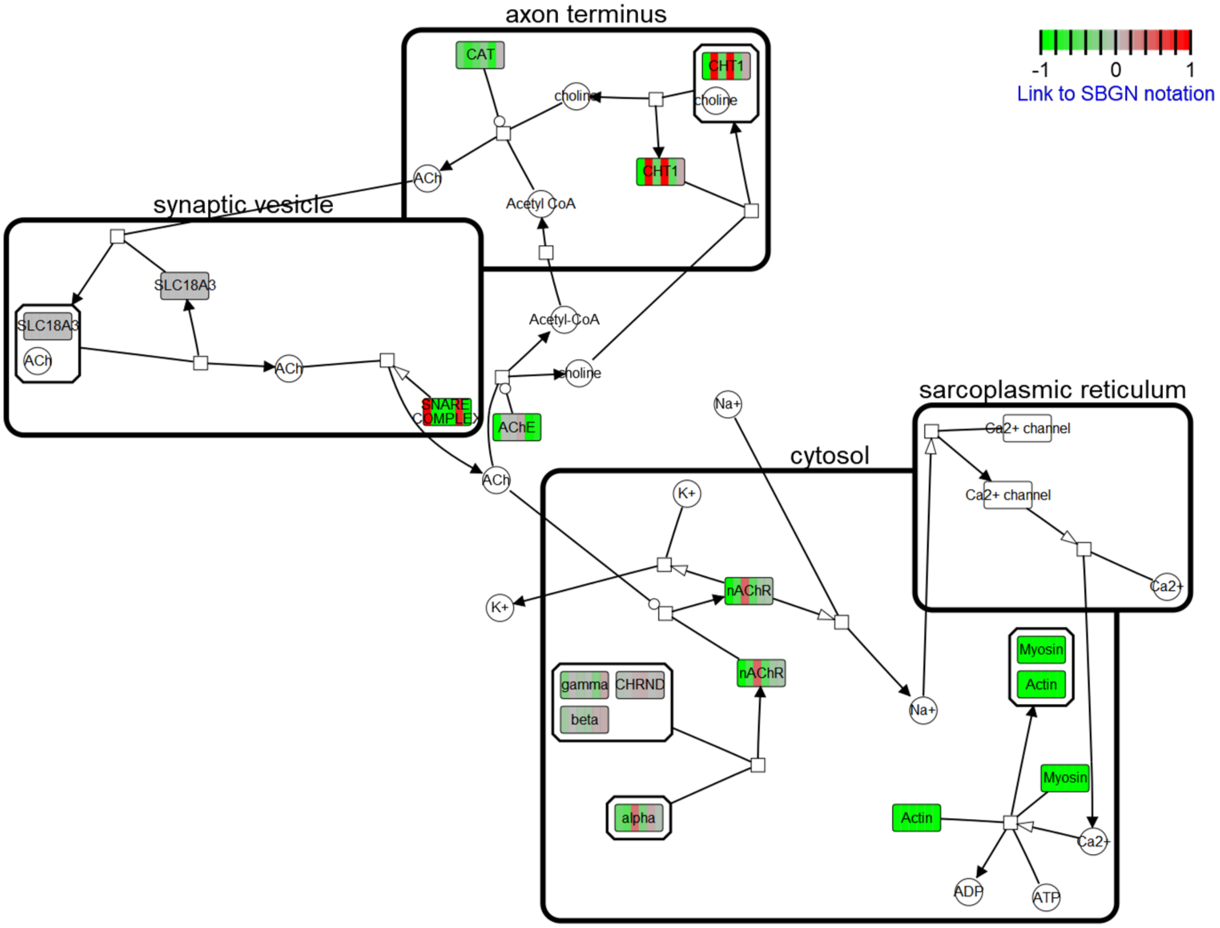


Supplementary Figure 6. Example SBGNview graph: Nicotinic acetylcholine receptor signaling pathway (PANTHER pathway P00044) down-regulated in breast cancer (GEO GSE16873).
